# Inverse Neural-Kinematic Dynamics Revealed by RQA During Motor Skill Learning

**DOI:** 10.1101/2025.11.03.685985

**Authors:** Navid Entezari, Zakieh Hassanzadeh, Fariba Bahrami, Ahmad Kalhor

**Author notes:** Contributing authors. These authors contributed equally to this work (Fariba Bahrami, Ahmad Kalhor).

## Abstract

Motor skill acquisition involves dynamic co-adaptation of brain and behavior, yet the temporal coupling between neural and kinematic processes during learning remains unclear. Traditional analyses often treat the brain and behavior independently, potentially overlooking their interactive dynamics. In this study, we applied Recurrence Quantification Analysis (RQA), a nonlinear method for assessing temporal structure in complex systems, to both EEG and foot trajectory data collected during a seven-session motor learning task. Twelve healthy adults practiced drawing five shapes with their dominant foot on a digital tablet while EEG was recorded from sensorimotor cortex electrodes. We found a fundamental inverse relationship between neural and behavioral dynamics: as motor performance improved, EEG determinism in the left hemisphere increased by 2.7% (p<0.001), while kinematic determinism decreased by 18.4% (p<0.001), indicating more organized brain activity supporting more flexible movement patterns. EEG laminarity also increased by 1.5% (p<0.001), while kinematic laminarity decreased by 15.2% (p<0.001). Shape-specific analysis revealed that patterns B and M elicited the most pronounced neural-behavioral adaptations. These findings suggest that increasingly structured neural dynamics facilitate adaptive motor execution. Our dual-domain RQA framework offers a powerful tool to quantify learning efficiency and uncover the dynamic coupling between neural reorganization and behavioral adaptation.

## Introduction

Motor learning involves the acquisition and refinement of skilled movements through repeated practice, underpinned by the dynamic interaction between neural processes and motor behavior. While classical EEG markers such as movement-related cortical potentials (MRCPs) and event-related desynchronization/synchronization (ERD/ERS) have deepened our understanding of motor function (Pfurtscheller & Da Silva, 1999; Jochumsen et al., 2017), these approaches often rely on linear, time-averaged features that obscure the rich temporal complexity of brain dynamics during learning (Cohen, 2014; Stam, 2005).

Electroencephalography (EEG), with its high temporal resolution, offers a window into real-time neural processing. However, traditional EEG analysis may overlook the nonlinear and evolving nature of motor-related neural activity. This limitation restricts our understanding of how the brain dynamically adapts in tandem with behavioral improvements over the course of learning (Stam, 2005).

Recent research has highlighted the central role of the sensorimotor cortex in motor skill acquisition. The sensorimotor cortex, a key region for planning and executing motor actions, plays a crucial role in skill acquisition by optimizing sensorimotor and cognitive parameters to achieve motor goals (Krakauer et al., 2019). During skill acquisition, distinct patterns of functional connectivity with the motor cortex emerge, reflecting various components of sensorimotor learning, including implicit processes, performance errors, and explicit strategy use. This highlights the sensorimotor cortex’s central role in the brain’s dynamic adaptation during motor skill acquisition (Areshenkoff et al., 2024). These findings suggest that understanding motor learning requires analytical approaches capable of capturing the complex, time-varying dynamics of neural connectivity patterns.

Nonlinear time-series methods such as Recurrence Quantification Analysis (RQA) provide a promising alternative for capturing these dynamic processes. RQA quantifies the recurrence structure of time-series data and captures features such as determinism, laminarity, and entropy—metrics sensitive to the stability, complexity, and transitions within dynamic systems. It has been effectively applied to characterize gait variability in pathological conditions (Mengarelli et al., 2021), assess functional brain connectivity in neuropsychiatric disorders (Jonak et al., 2020), and analyze cortical dynamics during motor execution (Pitsik et al., 2020). RQA is particularly well-suited for investigating motor learning because it can detect subtle changes in the temporal organization of neural signals that may precede or accompany behavioral improvements, providing insights into the mechanisms underlying motor plasticity. Despite these applications, a critical gap remains in understanding how recurrence patterns in neural activity relate to behavioral learning trajectories over extended practice periods, as traditional approaches analyze neural and behavioral data separately, potentially missing the coupled dynamics that characterize skill acquisition.

In this study, we address this gap by applying RQA to EEG signals recorded from the sensorimotor cortex and to foot-based kinematic data collected during the learning of a novel shape-drawing task. Participants completed seven sessions of training, enabling us to track how recurrence-based features evolve across time. Our aim is to capture the co-adaptive dynamics between brain and behavior during motor skill acquisition, and to identify patterns of neural plasticity and movement flexibility that emerge with practice.

## Methods

### Dataset

This study analyzed data from experiments originally conducted at the Human Motor Control and Computational Neuroscience Laboratory at the University of Tehran and the National Brain Mapping Laboratory. The experimental protocol was approved by the Research Ethics Committee of Iran University of Medical Sciences. The same dataset was previously analyzed in Khosravi et al. (2023).

Twelve healthy volunteers (6 males, 6 females; mean age 23±2 years) participated in the original study. All were right-foot dominant with no history of neurological conditions. Participants completed seven sessions over two weeks, learning to draw five geometric patterns (B, M, O, Star, Clover) with their dominant foot using a pen secured to a digital tablet (Wacom), with visual feedback displayed on screen. To maintain task consistency, participants were instructed to prioritize both speed and accuracy and were not permitted to look at their feet. Rest periods and task cues were displayed on the screen (**Figure 1**).

**Figure 1.**
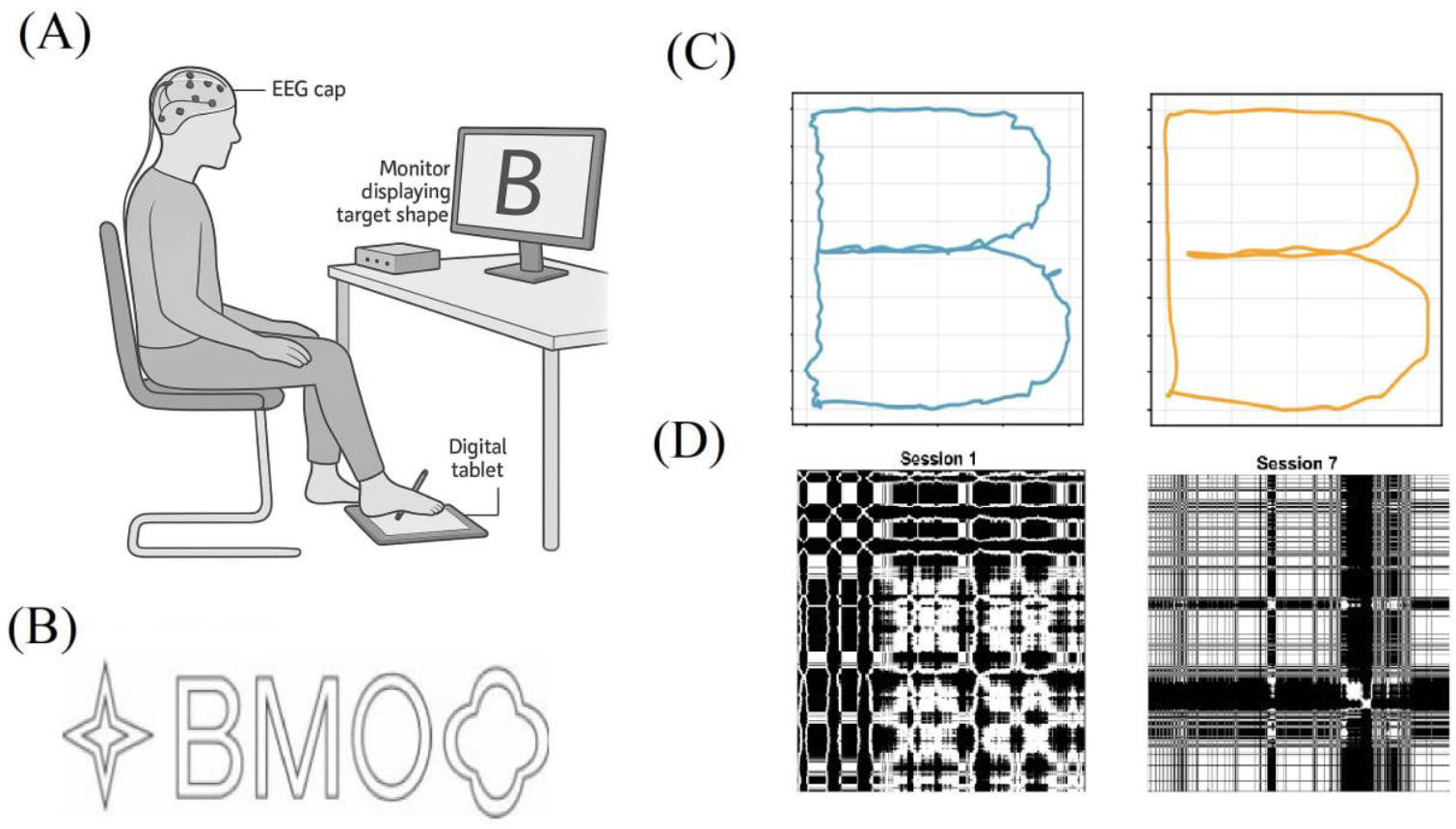
Experimental Setup and Task Design. (A) Participant positioning during the foot-based drawing task, showing the seated position with the dominant foot controlling a pen on a digital tablet while EEG is recorded. (B) Target shapes that participants were instructed to draw: a four-pointed star, the letter “B”, the letter “M”, the letter “O”, and a clover shape. (C) Representative kinematic trajectories showing motor learning progression for shape “B” from early learning to advanced learning, demonstrating improved accuracy and smoothness over the training period. (D) Recurrence plots illustrating the evolution of movement dynamics across learning sessions, with Session 1 (left) showing highly structured, deterministic patterns and Session 7 (right) displaying more fragmented, variable recurrence structures, reflecting the transition from rigid to flexible movement control during skill acquisition.

The anonymized dataset used in the present analysis was accessed by the authors between 01/05/2023 and 30/05/2025. All data were fully anonymized before being shared with the authors.

### Data Preprocessing

#### EEG Preprocessing

The preprocessing pipeline involved: (1) removal of EOG and synchronization channels; (2) removal of 100-second resting periods between trials; (3) visual inspection to identify and remove artifacts using EEGLAB’s artifact rejection tools; (4) bandpass filtering (0.5-45 Hz) to remove low-frequency drifts and high-frequency noise; (5) Independent Component Analysis (ICA) using the Infomax algorithm to remove artifact components related to eye movements and muscle activity. Approximately 30% of independent components were removed based on spatial, temporal, and spectral characteristics. All processing was performed using MATLAB and EEGLAB (Delorme & Makeig, 2004).

#### Kinematic Preprocessing

Drawing-related segments were extracted using synchronization signals to identify start and end points of each pattern. To eliminate foot tremor, a 10th-order Butterworth low-pass filter with 5 Hz cutoff was applied. Velocity of foot movement was calculated in two perpendicular directions using pen position data.

### Regions of Interest

The sensorimotor cortex plays a critical role in motor learning, with primary motor cortex and associated areas showing structural and functional changes during skill acquisition (Krakauer et al., 2019). The primary motor cortex is the main contributor to generating neural impulses that control movement execution, requiring the least amount of electrical current to elicit movements compared to other motor areas (Yip et al., 2024; Knierim, 2020). To investigate these functional changes during motor learning, EEG signals were analyzed from nine electrodes (Cp4, C4, Fc4, Cpz, Cz, Fcz, Cp3, C3, Fc3) associated with the sensorimotor region. Based on their anatomical locations, these electrodes were grouped into three areas:

1. right hemisphere: Cp3, C3, Fc3
2. left hemisphere: Cp4, C4, Fc4
3. longitudinal fissure: Cpz, Cz, Fcz

This grouping approach follows previous work that demonstrated the sensitivity of recurrence-based features to changes in motor-related EEG activity (Pitsik et al., 2020). The focus on sensorimotor areas is particularly relevant for foot-drawing tasks, as motor acuity improvements during novel skill acquisition are accompanied by neuroplastic changes in primary and premotor cortex (Krakauer et al., 2019).

### Recurrence quantification analysis

To quantify the nonlinear dynamics of neural and movement data, we implemented recurrence quantification analysis (RQA). This approach characterizes complex time series by analyzing recurrence patterns in the system’s state space through recurrence plots (RPs), which visualize when a dynamical system revisits similar states (Eckmann et al., 1987).

#### Phase Space Reconstruction

Before creating recurrence plots, we reconstructed the phase space of our time series using time-delay embedding. This technique transforms a single variable time series x(t) into a multidimensional state space representation X(t) according to:

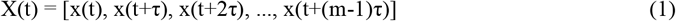

where τ is the time delay and m is the embedding dimension. These parameters determine how effectively the system’s dynamics are captured: τ controls the temporal separation between successive coordinates, while m defines the dimensionality of the reconstructed space.

For each EEG epoch and kinematic data segment, we determined optimal embedding parameters using established methods. The embedding dimension m was calculated using the false nearest neighbors method, which identifies the minimum dimension needed to unfold the attractor. The time delay τ was determined using the mutual information method, which identifies delays that minimize redundancy between coordinates. Given the multidimensional nature of our datasets, we applied the approach proposed by Wallot and Mønster (2018) for robust parameter estimation in multidimensional recurrence quantification analysis (Wallot et al., 2016). Since embedding parameter values varied slightly across epochs, we used the median values from our dataset (m = 4 and τ = 13) for consistency across all analyses.

#### Recurrence Plot Construction

After phase space reconstruction, recurrence plots were constructed using the Cross Recurrence Plot MATLAB toolbox (Marwan et al.,2007). The recurrence plot was constructed according to:

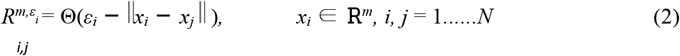

where xi and xj are state vectors in the m-dimensional reconstructed phase space, N represents the number of state vectors after phase space reconstruction, Θ is the Heaviside function, and ε is the threshold parameter.

Selecting an appropriate threshold is critical for revealing meaningful recurrence structures. Too small a threshold yields few recurrences and sparse patterns, while too large a threshold creates excessive recurrences that might obscure the system’s dynamics. For EEG data, we maintained recurrence rates below 10% to avoid spurious correlations, following recommendations by Mindlin and Gilmore (1992) for neural data analysis. For kinematic data, which typically exhibits greater deterministic structure, we allowed recurrence rates up to 20%.

We systematically varied ε from 0.2 to 0.8 in steps of 0.05 for EEG analysis, and from 0.05 to 0.7 in steps of 0.05 for kinematic data. This approach allowed us to identify the most sensitive threshold values for detecting learning-related changes. Final analyses were conducted using ε = 0.65 for EEG and ε = 0.7 for kinematic data. RQA measures were calculated using a sliding window approach with 1,000 samples per window and 50% overlap (500 samples) between successive windows. This windowing strategy enables the assessment of dynamic changes over time while maintaining sufficient temporal resolution. The computed RQA measures from all windows were then averaged to yield stable estimates for each experimental condition.

#### RQA Measures

We focused on four key RQA measures to characterize the temporal structure of both EEG and kinematic signals:

##### Determinism (DET)

The proportion of recurrence points that form diagonal lines of at least length *l*_*min*_ indicating predictable and repeated patterns in the signal.

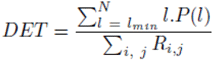

where P(l) is the histogram of diagonal line lengths and *R*_*i, j*_ is the recurrence matrix.

##### Diagonal Line Entropy

Measures the complexity of the system by quantifying the distribution of diagonal line lengths.

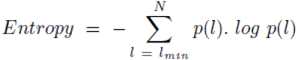

where 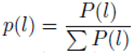 is the probability of diagonal line length l.

##### Laminarity (LAM)

The fraction of recurrence points forming vertical lines of at least length *v*_*min*_, often interpreted as reflecting intermittent stable or laminar phases.

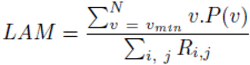

where P(v) is the histogram of vertical line lengths.

##### Mean Diagonal Line Length (L)

Represents the average length of all diagonal lines and is related to the predictability of the system.)

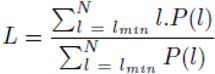

### Statistical Analysis

RQA measures were calculated for both EEG signals and kinematic data from three sessions (first, fourth, and seventh) to track changes across the learning period. To examine these changes, we used Repeated Measures ANOVA with statistical tests performed for optimal epsilon values satisfying sphericity and normality conditions. We analyzed three types of effects: (1) main effect of session, indicating learning-related changes over time; (2) main effect of shape, showing differences between shapes; and (3) interaction effects between shape-specific learning rates. Post-hoc comparisons with Bonferroni correction were conducted for significant effects. Statistical significance was set at p < 0.05.

## Results

We analyzed recurrence quantification measures from both kinematic data and EEG signals across three time points to track changes during motor learning. Our analysis focused on main effects of session, differences between shapes, and interaction effects (shape-specific learning trajectories).

### Motor Learning Progression

After 7 sessions of practice, improvement in performance was evident across all participants. **Figure 2** demonstrates the motor learning progression across three sessions for all five geometric shapes. The kinematic trajectories show clear improvements in movement accuracy and smoothness over the learning period. In Session 1, participants’ movements were characterized by irregular, jerky trajectories with frequent corrections and deviations from the target shapes. By Session 4, movements became more controlled and closer to the intended geometric forms, though some variability remained. Session 7 trajectories demonstrated marked improvement, with smoother, more precise movements that closely approximated the target shapes. The learning progression was particularly evident in complex shapes such as the star and clover patterns, where initial attempts showed significant spatial errors. As training progressed, participants developed more efficient motor strategies, resulting in reduced movement variability and improved spatial accuracy.

**Figure 2.**
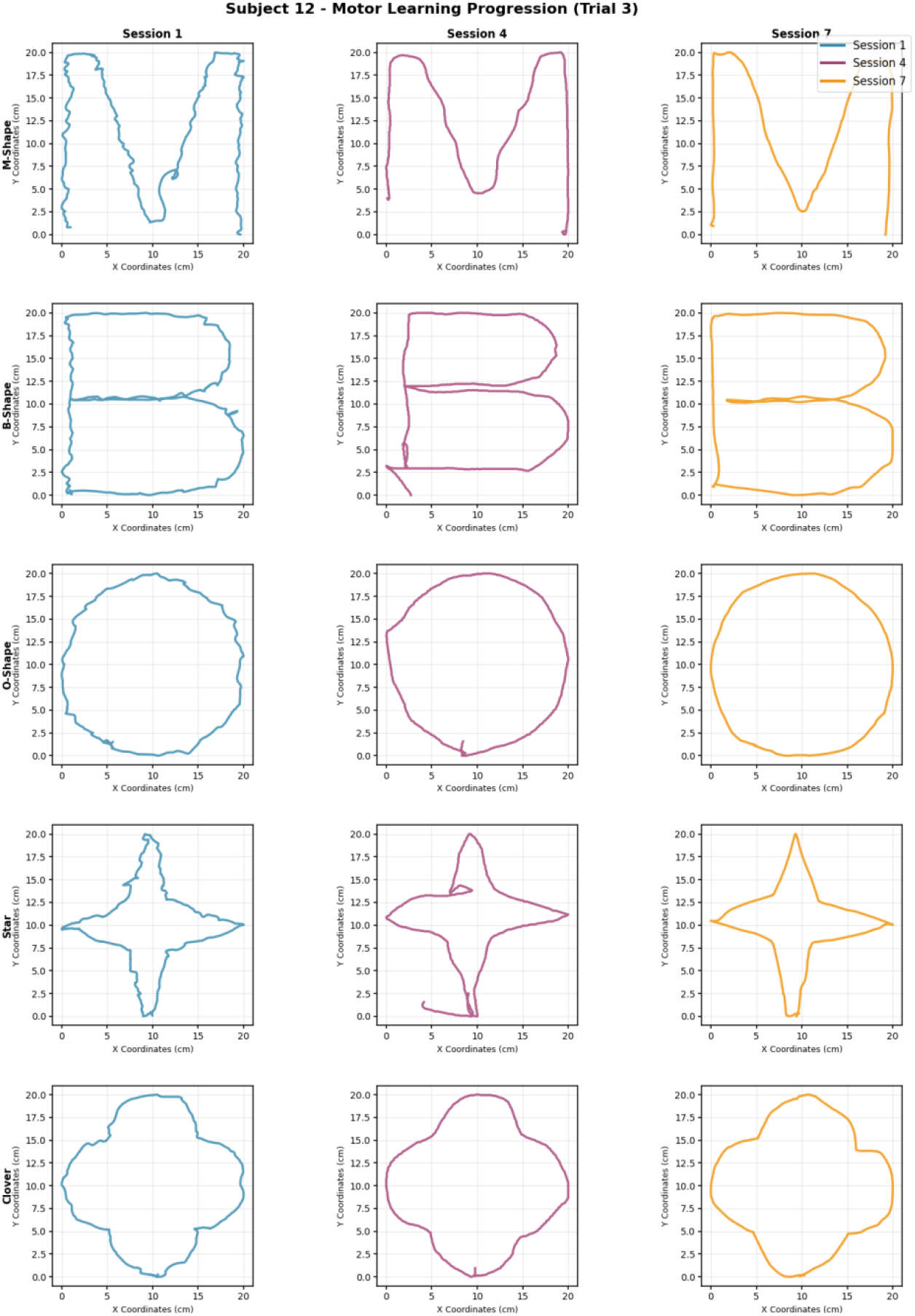
Motor learning progression across training sessions. Kinematic trajectories for all five geometric shapes drawn by a representative participant during Session 1 (blue), Session 4 (purple), and Session 7 (orange), demonstrating systematic improvements in movement accuracy and smoothness over the learning period.

### Kinematic RQA Findings

#### Changes in Recurrence Patterns

Figure 3 presents kinematic recurrence plots showing velocity data across learning sessions for a representative participant performing shape M. Visual analysis of recurrence plots examines global patterns (typology) and local structures (textures) that reveal system dynamics (Marwan et al., 2007). The plots demonstrate systematic changes in movement dynamics throughout the learning process: Session 1 exhibits highly structured, deterministic recurrence patterns with prominent diagonal lines and grid-like organization, reflecting rigid and predictable movement velocities characteristic of early motor learning. Session 4 shows intermediate organization with mixed structural elements, indicating the emergence of movement variability. Session 7 displays fragmented recurrence patterns with reduced diagonal structures and shorter segments, representing the development of more flexible and variable movement dynamics. This visual progression from organized to disrupted structure provides intuitive validation that motor skill acquisition involves a transition from rigid to flexible movement patterns.

**Figure 3.**
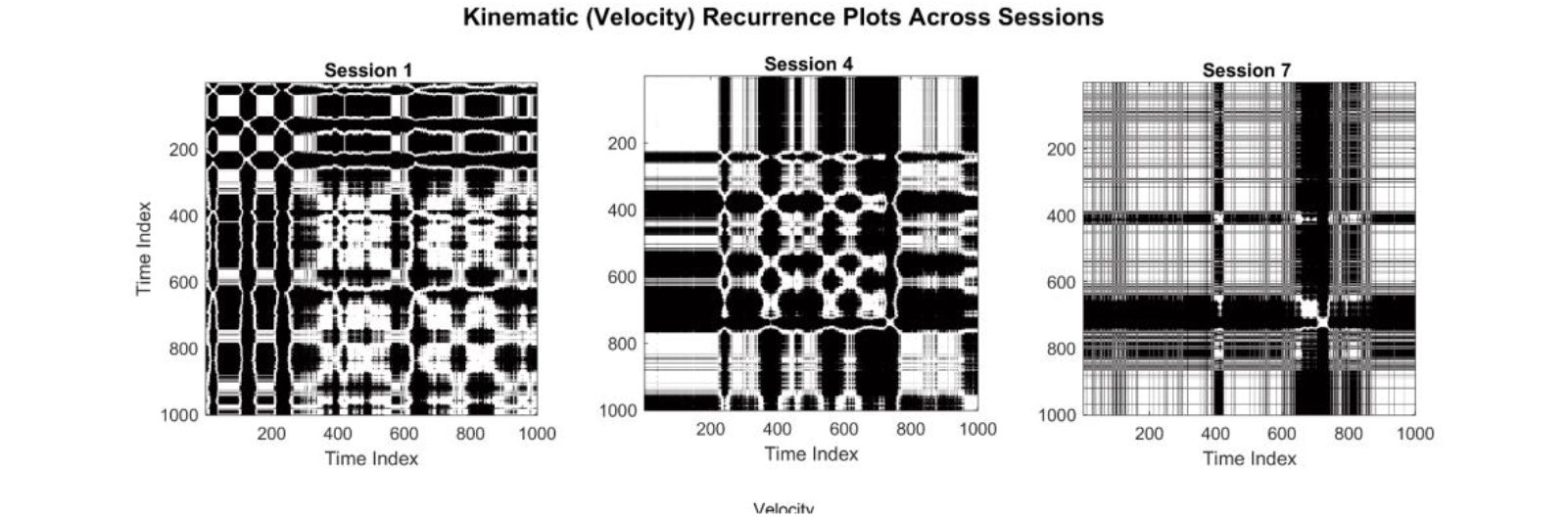
Kinematic recurrence plots showing velocity data across learning sessions for a representative participant. Note the progressive changes in recurrence structure from Session 1 (left) to Session 7 (right), demonstrating the evolution from highly structured, deterministic movement patterns to more variable, less predictable dynamics. The reduction in determinism and laminarity quantified by RQA metrics reflects this transition toward more flexible and adaptive motor control strategies during learning.

#### Main Effects of Learning

The RQA analysis of behavioral kinematic data revealed consistent decreases across all measures during motor learning, with statistically significant reductions from Day 1 to Day 7 (**Figure 4**). The most reduction was observed in Mean Diagonal Line Length (-45.4%, p < 0.001), followed by Diagonal Line Entropy (- 29.2%, p < 0.001), Determinism (-18.4%, p < 0.001), and Laminarity (-15.2%, p < 0.001). These decreases reflect the emergence of more variable and less predictable movement patterns as participants developed motor expertise, consistent with theories of skill acquisition that emphasize the transition from rigid, highly structured movements to more flexible and adaptive motor control.

**Figure 4.**
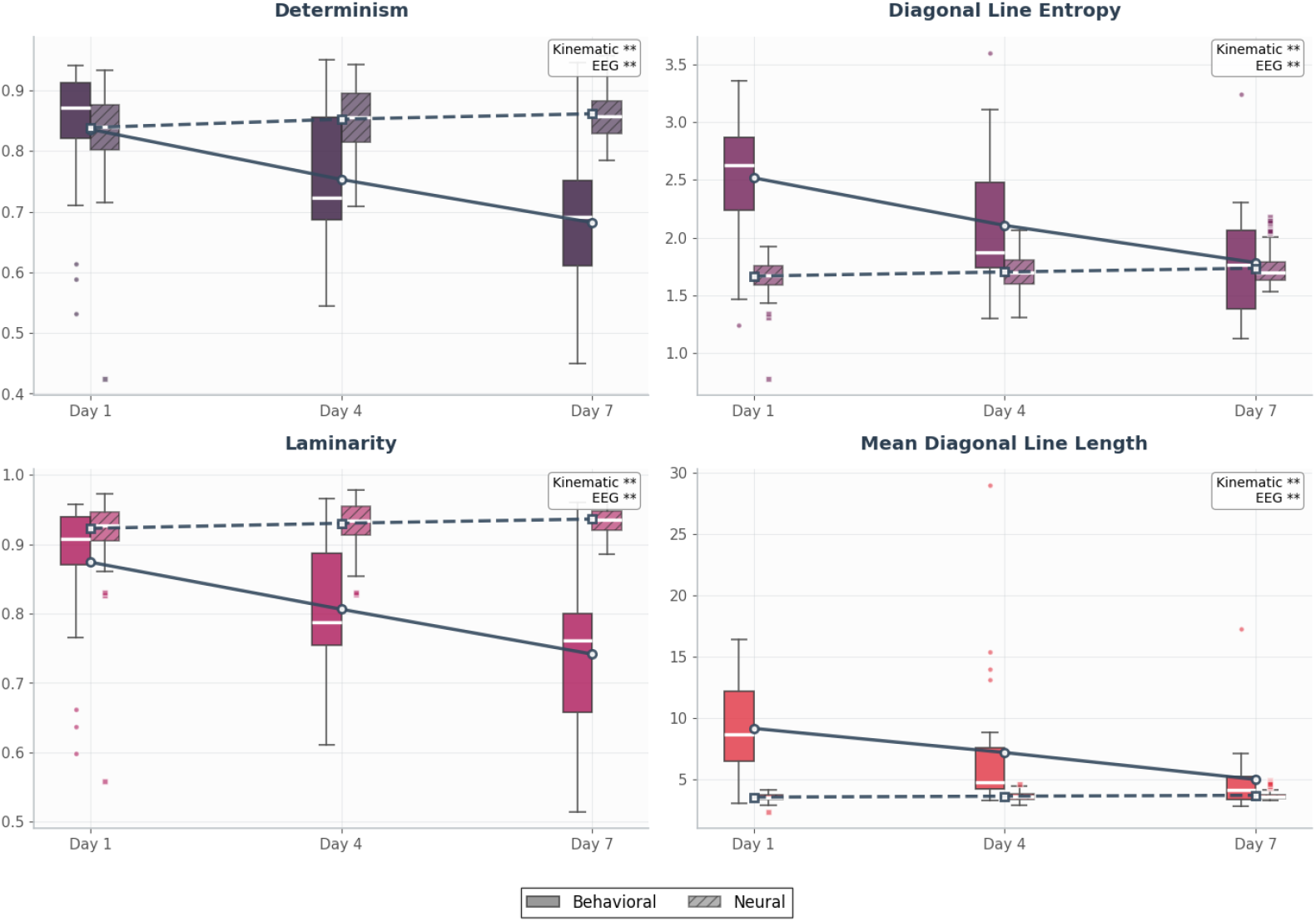
behavioral and neural RQA dynamics during motor learning. Box plots show four RQA metrics across training sessions (Day 1, Day 4, Day 7) for kinematic (solid boxes) and EEG (hatched boxes) data (left hemisphere). Significance indicators denote Day 1 vs Day 7 comparisons (** = p < 0.01). All behavioral measures decreased significantly (15-45% reductions, all p < 0.001), while neural measures increased modestly but significantly (1.5-4.2% increases, all p < 0.001), revealing opposing patterns of dynamical reorganization during skill acquisition.

#### Shape-Specific Learning Effects

While RQA measures did not reveal significant differences between shapes in general, shape-specific learning trajectories (session × shape interaction) showed distinct patterns (**Figure 6**). Significant decreases in determinism were observed for Shape B (p = 0.009), Shape Clover (p = 0.006), and Shape M (p = 0.010). Shape O showed a moderate decrease (p = 0.029), while Shape Star did not reach statistical significance (p = 0.164). For diagonal line entropy, significant reductions were observed for Shape B (p = 0.011), Shape Clover (p = 0.010), and Shape M (p = 0.008), with Shape O showing a milder but still significant change (p = 0.025). Shape Star did not show a significant trend (p = 0.147).

Laminarity analysis revealed significant decreases for Shape B (p = 0.008), Shape Clover (p = 0.006), and Shape M (p = 0.008). Shape O also exhibited a significant reduction (p = 0.028), while changes in Shape Star were not significant (p = 0.179). Changes in mean diagonal line length were more variable across shapes. Significant decreases were observed for Shape B (p = 0.026), Shape Clover (p = 0.017), and Shape O (p = 0.033). Shape M approached significance (p = 0.061), while Shape Star showed no significant change (p = 0.396). These patterns suggest that Shapes B, Clover, and M facilitated the most consistent motor learning across all measures, while Shape Star showed the least learning-related changes.

### EEG RQA Findings

#### Hemisphere-Specific Neural Adaptations

Analysis of EEG signals from the sensorimotor cortex revealed significant learning-related changes exclusively in the left hemisphere (electrodes Cp3, C3, Fc3), while the right hemisphere (Cp4, C4, Fc4) and midline areas (Cpz, Cz, Fcz) showed no significant adaptations. This lateralization pattern is particularly notable given that participants performed the task with their dominant (right) foot, suggesting that the contralateral left hemisphere undergoes specific reorganization during novel motor skill acquisition.

Figure 5 presents EEG recurrence plots from all three sensorimotor regions (left hemisphere, right hemisphere, and midline) across the three learning sessions (Sessions 1, 4, and 7) for Shape B. The 9 recurrence plots reveal regional and temporal differences in neural dynamics throughout the learning process. The left hemisphere plots show significant changes from Session 1 to Session 7, with quantitative analysis revealing significant increases in determinism (2.7% increase, p < 0.001) and laminarity (1.5% increase, p < 0.001) as detailed below. In contrast, the right hemisphere and midline regions show minimal changes across sessions, highlighting the hemisphere-specific nature of neural adaptation during this motor learning task.

**Figure 5.**
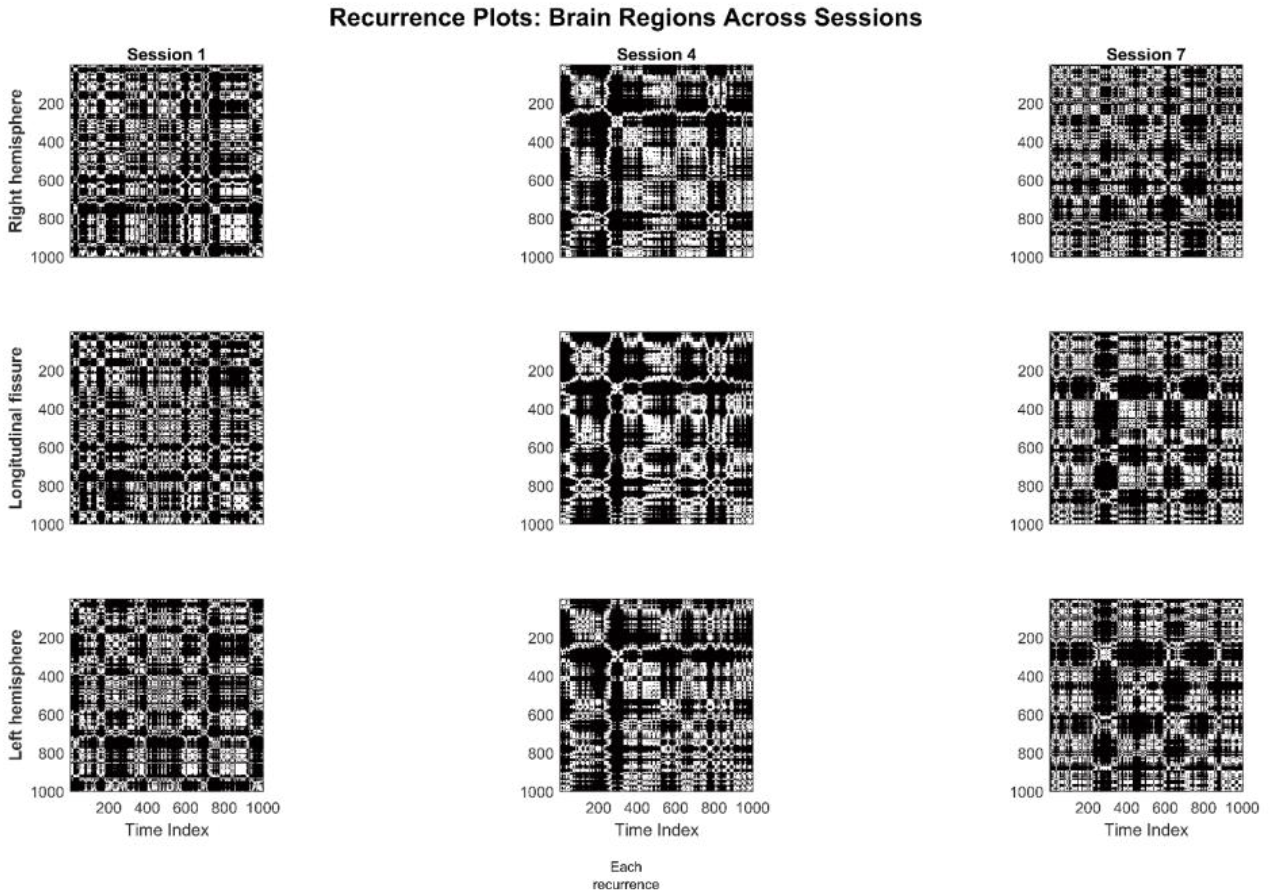
EEG recurrence plots from three sensorimotor regions (left, right, midline) across Sessions 1, 4, and 7 for Shape B. The nine plots reveal regional differences in neural dynamics, with the left hemisphere (bottom row) showing progressive organization from sparse, fragmented patterns to structured, coherent diagonal formations, while right hemisphere and midline regions display minimal changes across sessions.

#### Main Effects of Neural Learning

Neural RQA measures derived from EEG data showed a distinctly different pattern compared to the behavioral findings, with modest but highly significant increases across all metrics from Day 1 to Day 7. Specifically, Mean Diagonal Line Length increased by 4.2% (p < 0.001), Diagonal Line Entropy by 4.1% (p < 0.001), Determinism by 2.7% (p < 0.001), and Laminarity by 1.5% (p < 0.001). This pattern suggests that the underlying neural control strategies become more structured and predictable with learning. The increased neural recurrence dynamics indicate that motor skill acquisition involves the development of more stable and organized neural control patterns, which may provide the foundation for the enhanced behavioral flexibility observed in the kinematic data.

#### Shape-Specific Neural Adaptations

Similar to the kinematic data, we found shape-specific learning effects (session × shape interaction) in the EEG data. **Figure 6** presents the learning-related changes in determinism, diagonal line entropy, and laminarity in the left hemisphere for each movement shape. Significant changes were observed for Shape B in both determinism (p = 0.039) and diagonal line entropy (p = 0.039). Additionally, Shape M showed a significant increase in diagonal line entropy (p = 0.085) and a marginal change in determinism (p = 0.134). For laminarity, no shape exhibited statistically significant changes at the threshold (p < 0.05), though Shape B approached significance (p = 0.078).

**Figure 6.**
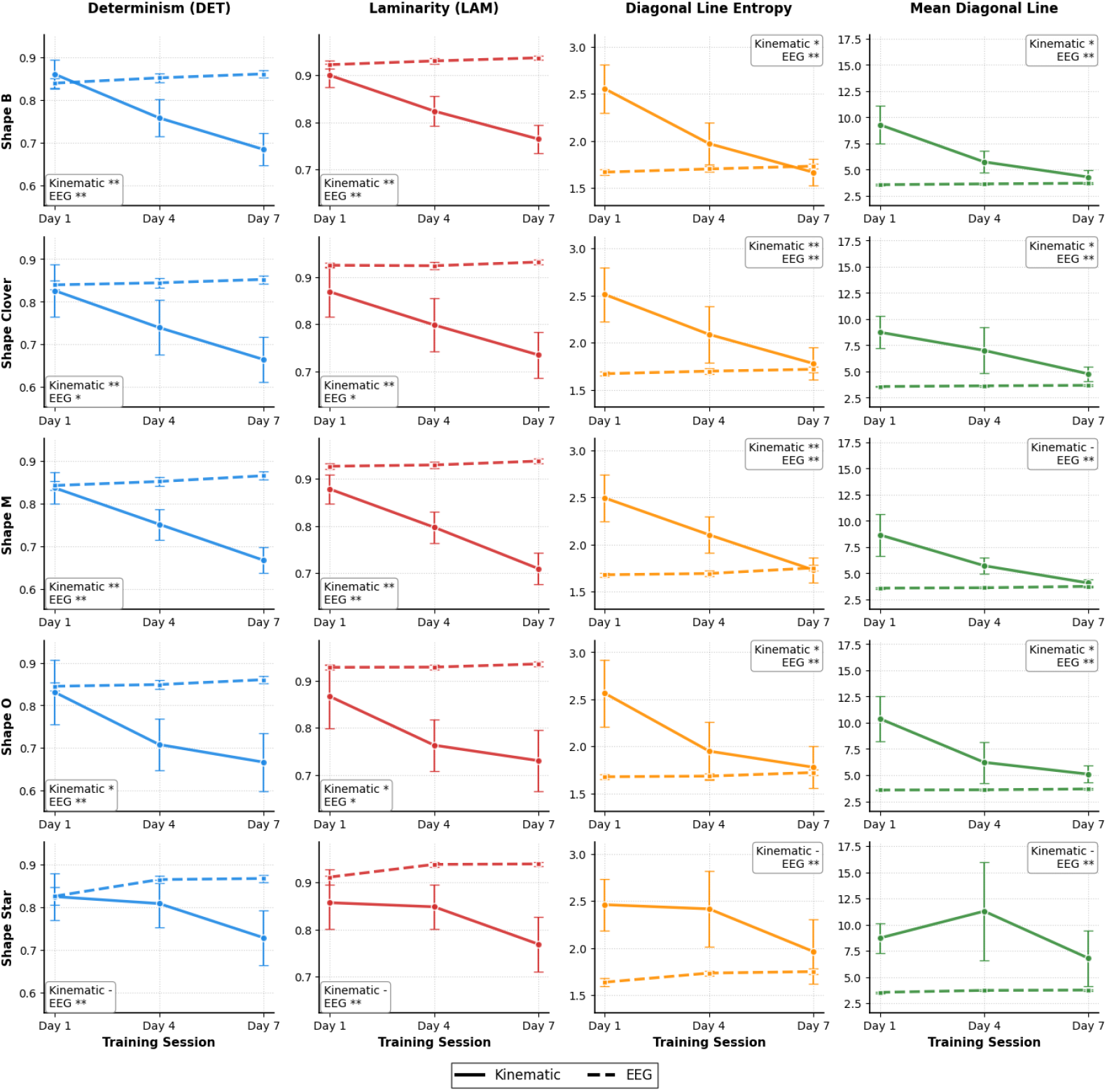
Shape-specific RQA dynamics during motor learning. RQA metrics (columns) across shape conditions (rows) for three training sessions. Kinematic data (solid lines, circles) and EEG data (dashed lines, squares) show shape-dependent learning patterns. Individual legends indicate Day 1 vs Day 7 significance (** = p < 0.01, * = p < 0.05, - = not significant). Error bars represent standard error of the mean.

#### Relationship Between Neural and Kinematic Changes

A key finding of our study is the inverse relationship between neural and kinematic determinism during skill acquisition. As participants improved in the task, we observed increased determinism in EEG signals (2.7% increase, p < 0.001) while simultaneously seeing decreased determinism in movement kinematics (18.4% decrease, p < 0.001). This pattern suggests a fundamental aspect of motor learning where more stable and efficient neural processing supports more flexible and adaptive movement patterns. Similarly, while laminarity (reflecting stable, low-variability phases) decreased in kinematic data (15.2% decrease, p < 0.001), it increased in EEG signals (1.5% increase, p < 0.001), further supporting this reciprocal relationship between neural and motor dynamics during learning.

## Discussion

The application of recurrence quantification analysis to both kinematic data and EEG signals reveals significant dynamic changes during motor learning. This dual-domain approach provides novel insights into how neural processing and movement patterns co-adapt during the acquisition of a novel foot-based drawing task.

### Neural and Kinematic Adaptations During Motor Learning

The decreasing determinism observed in participants’ foot movement velocity during training indicates increased movement flexibility and adaptability. This shift from more rigid, predictable movement patterns to more variable, context-sensitive movements aligns with established theories of motor learning, where early learning phases are characterized by stereotyped movements that gradually become more fluid and efficient with practice. The decrease in laminarity suggests improved neuromuscular coordination and faster adjustments in movement execution, reflecting the development of more efficient internal models and reduced reliance on real-time sensory feedback corrections resulting from practice (Shumway-Cook & Woollacott, 2007).

Notably, the reduction in mean diagonal line length indicates that participants follow specific movement patterns for shorter durations, suggesting the emergence of more automated, feedforward motor programs with increased fluidity and adaptability to task demands. The observed decrease in diagonal line entropy points to increased uniformity of predictable structures within movements, aligning with theoretical models suggesting that efficient motor programs reduce unnecessary complexity (Schmidt et al., 2018), focusing on task-relevant movement patterns while eliminating extraneous elements. These RQA-based findings provide novel insights into the temporal dynamics of motor skill acquisition, as this study represents one of the first applications of mean diagonal line length and diagonal line entropy to characterize neural-behavioral coupling during motor learning.

Concurrent neural adaptations in the left hemisphere sensorimotor cortex (Cp3, C3, Fc3) provide evidence of neural reorganization during motor learning. The increase in neural determinism suggests that brain activity becomes more structured and predictable as skill improves, likely reflecting the formation of stable cortical patterns related to improved motor planning and execution (Azim & Seki, 2019; Franklin et al., 2016). This interpretation is supported by Bonnette et al. (2024), who demonstrated that higher neural determinism in sensorimotor regions correlates with better motor control and reduced injury risk during complex movements, suggesting that increased neural organization reflects enhanced motor function. Our finding of significant changes primarily in the left hemisphere contributes to understanding hemispheric specialization in motor learning. While Bonnette et al. observed these patterns during acute movement phases, our study extends this principle to demonstrate how neural determinism increases over multiple learning sessions, suggesting that learning-related neural reorganization builds upon previous work showing that motor execution affects sensorimotor cortex signal complexity (Pitsik et al., 2020). While immediate execution produces more regular neural dynamics, our study reveals how these patterns strengthen over multiple learning sessions, suggesting both immediate task-specific synchronization and longer-term efficiency-driven reorganization.

The modest increases (2-3%) in EEG determinism and laminarity observed in the sensorimotor cortex electrodes (Cp4, C4, Fc4, Cpz, Cz, Fcz, Cp3, C3, Fc3) reflect the inherent characteristics of these regions during motor learning. The sensorimotor cortex is among the most highly organized and structured brain regions even before learning begins which shows maximal synchronization during active movement execution (Krakauer et al., 2019). Unlike other cortical regions that undergo dramatic reorganization during learning, the sensorimotor cortex shows less reorganization since it primarily functions as the final common pathway for motor execution rather than the primary locus of motor learning plasticity. The major learning-related plasticity during skill acquisition occurs predominantly in areas such as the cerebellum for error correction and fine-tuning, basal ganglia for action selection and habit formation, and premotor areas for motor planning and sequencing (Krakauer et al., 2019).

Since the sensorimotor cortex is already operating near optimal efficiency levels during motor tasks, there is limited capacity for further increases in neural organization - a phenomenon analogous to ceiling effects in motor learning (Hardwick et al., 2013). The high initial values (determinism ∼84%, laminarity ∼92%) in both kinematic and neural RQA measures reflect characteristics of the cognitive stage of motor learning, where novice learners exhibit highly structured, rigid movement patterns as they consciously attempt to control this novel foot-drawing task (Krakauer et al., 2019). During this early learning phase, the sensorimotor cortex must work intensively to coordinate the unfamiliar motor demands, resulting in highly organized neural firing patterns that support the stereotyped, effortful movements characteristic of novice performance. The small but consistent changes we observed likely represent fine-tuning of existing sensorimotor maps and subtle optimization of neural firing patterns specific to the foot-drawing task (statistically significant effects), rather than wholesale cortical reorganization (low magnitude changes) (Hardwick et al., 2013). The increase in neural laminarity suggests the emergence of more stable, low-variability neural states, which may indicate reduced cognitive load during task performance. These states likely reflect the early consolidation of task-specific neural representations, enabling more automatic execution with decreased reliance on attentional control.

Analysis of shape-specific learning trajectories revealed intriguing differences in adaptation patterns. Drawing the star shape appeared most challenging, showing the least change in both neural and kinematic metrics, suggesting that complex shapes with sharp angles and multiple direction changes may require longer practice periods or different learning strategies. In contrast, patterns B and M showed the most pronounced changes in both domains, suggesting these shapes may be particularly suitable for rehabilitation applications due to their optimal challenge level that promotes neuromotor adaptation without overwhelming the learner’s capacity.

### Inverse Coupling and Neuromuscular Coordination Implications

One of the most striking findings is the inverse relationship between neural and kinematic determinism during motor skill acquisition. As participants became more proficient, neural determinism increased while kinematic determinism decreased, suggesting a fundamental principle in sensorimotor learning: more stable and efficient neural processing supports more flexible and adaptive movement execution. This inverse relationship can be understood through neuromotor efficiency—as learning progresses, the brain develops more consistent neural patterns that require less variable top-down control, enabling greater flexibility in movement execution and adaptive responses to task demands.

This pattern contrasts with pathological conditions, where increased neural determinism often reflects constrained adaptability. Patients with neurological disorders exhibit abnormally high EEG determinism, indicating less flexible neural processing in multiple sclerosis, Alzheimer’s disease, Parkinson’s disease, and epilepsy (Carrubba et al., 2012, 2019). Conversely, vestibular dysfunction shows abnormally low kinematic determinism during walking, indicating chaotic movement patterns (Sylos Labini et al., 2012). These contrasting manifestations highlight that optimal sensorimotor function requires appropriate levels of determinism across physiological levels, with learning representing adaptive optimization and pathologies representing distinct forms of dysregulation.

The methodological framework established here opens possibilities for extending our approach to directly quantify temporal coupling between neural and behavioral signals. Cross-Recurrence Quantification Analysis (CRQA) and other nonlinear coupling techniques could assess coordination between multidimensional time-series, making them particularly suitable for analyzing complex neuromuscular coordination dynamics (Wallot et al., 2018; Wallot et al., 2019). Future research could apply these methods to simultaneously recorded EEG and EMG data during motor learning to quantify how neural-muscular coupling patterns evolve with skill acquisition, potentially providing more sensitive biomarkers for rehabilitation progress and motor recovery (Wang et al., 2024). Additionally, future studies should expand electrode coverage beyond the sensorimotor cortex to include cerebellar and subcortical regions using advanced source localization techniques, enabling investigation of broader neural network dynamics during motor learning. This expansion would capture not only proprioceptive processing but also visual-motor integration areas (occipital-parietal regions) and spatial learning circuits (frontal-parietal networks), allowing investigation of sensory reweighting dynamics (Assländer & Peterka, 2014) during skill acquisition.

